# Senescence targeting re-enables injury-responsive repair in a human RPE aging model

**DOI:** 10.64898/2026.06.26.722664

**Authors:** Birgit Ritschka, Clemens Etl, Fernando Becerril Perez, Keisuke Ishihara, Seba Almedawar, Marco Leyva Gonzalez, Heike Neubauer, Remko A. Bakker, Elly M. Tanaka

## Abstract

Aging is associated with progressive tissue dysfunction and impaired repair after injury. In the retinal pigment epithelium (RPE), these changes contribute to age-related macular degeneration (AMD), yet the mechanisms limiting repair remain incompletely understood. Here, we establish a longitudinal human embryonic stem cell (hESC)-derived RPE aging model that recapitulates key features of aged human donor RPE and combine it with mosaic cell ablation to assess injury-responsive repair. Although aged RPE cells initiate DNA synthesis after injury, they exhibit impaired mitotic progression, uncoupling S-phase entry from epithelial repopulation. Transcriptomic profiling links this defect to a senescence-associated program marked by inflammatory signaling and suppressed mitotic networks. Pharmacologic reduction of senescent cells with Navitoclax shifts aged RPE toward a younger transcriptional profile but does not induce repopulation by itself. Instead, senolytic treatment primes aged RPE for repair, improving epithelial density and homeostatic function only in response to injury, a strategy we term “senolytic priming.” These findings establish a human stem-cell-derived platform for investigating age-associated epithelial repair failure and show that aged human RPE retains latent repair capacity that can be re-enabled by targeting cellular senescence.

## INTRODUCTION

The human retina undergoes structural and functional changes during aging, contributing to declining visual function. A critical component of retinal health is the retinal pigment epithelium (RPE), a specialized monolayer that supports photoreceptors and maintains retinal homeostasis. RPE dysfunction contributes to age-related macular degeneration (AMD), a leading cause of irreversible central vision loss affecting ∼200 million people worldwide^1^. While anti-VEGF therapies have improved outcomes for neovascular AMD^2^, no effective treatments replace lost RPE cells or prevent tissue loss in atrophic AMD, which accounts for most cases.

A central unresolved question is whether human RPE retains meaningful repair capacity, and whether aging extinguishes this potential or constrains its execution. Although adult RPE is largely quiescent, *in vivo* and *in vitro* studies have documented limited compensatory responses to cell loss, including hypertrophy, multinucleation, and context-dependent cell-cycle re-entry, generally interpreted as non-productive or region-restricted attempts at repair^3–7^. Adult human RPE can also proliferate *ex vivo* under defined growth-factor conditions^8^. Nevertheless, intact tissue undergoes progressive cell loss with age^9,10^, compensated by hypertrophy^10,11^ and accompanied by barrier and phagocytic deficits^12,13^. These observations suggest that aged RPE may retain latent repair programs constrained by aging-associated changes. Testing this requires models that recapitulate human RPE aging while enabling mechanistic interrogation of injury responses.

Multiple processes have been implicated in age-related RPE dysfunction, including metabolic dysregulation, chronic inflammation, oxidative stress, complement activation, and cellular senescence^14–16^. Cellular senescence is a compelling candidate mechanism because it links stable cell-cycle arrest with inflammatory and tissue-remodeling signals^17,18^. Although senescence contributes to developmental remodeling and acute repair, persistent accumulation is associated with degenerative pathology^18,19^. RPE from AMD donor eyes exhibit senescence-associated features, including increased expression of cell-cycle inhibitors, senescence-associated β-galactosidase (SA-β-Gal) activity, and pro-inflammatory senescence-associated secretory phenotype (SASP) factors^20–22^. Senolytic agents that eliminate senescent cells have emerged as therapeutic candidates^23,24^ and have shown benefit in retinal disease models^25–27^. Whether senolytics improve RPE-intrinsic repair competence or act through indirect microenvironmental effects remains unknown, as does whether senescence directly constrains injury-responsive repair in human RPE.

To address these questions, we established a longitudinal hESC-derived model of RPE aging combined with a mosaic cell-ablation assay to interrogate injury-responsive repair. In this system, aged RPE re-entered S-phase after injury but failed to progress efficiently through mitosis, uncoupling DNA synthesis from productive epithelial repopulation. Transcriptomic profiling linked this defect to a senescence-associated program and suppression of mitotic genes. Senolytic intervention reduced senescent cell burden and shifted aged cultures toward a younger transcriptional state but did not induce repopulation. Instead, senolytic treatment enabled recovery of epithelial density and homeostatic function only when combined with injury cues, a framework we term “senolytic priming.” Together, these findings identify cellular senescence as a targetable barrier to epithelial repair competence and establish senolytic priming as a strategy for restoring repair in aged human tissues.

## RESULTS

### hESC-derived RPE cultures recapitulate aging-associated phenotypes over time *in vitro*

Limited access to human retinal tissue and the difficulty of longitudinally interrogating aging *in vivo* constrain mechanistic studies of RPE aging. To model progressive age-associated decline in a tractable human system, we established a longitudinal RPE culture platform using the H9 hESC line. hESCs were differentiated via neuroepithelial cyst formation^28,29^, matured as polarized monolayers on transwell inserts, and assessed from day 14 (D14) to D66 (Figure 1A). Based on functional and morphometric changes across this time course (Figures 1B, 1C, S1A, and S1B), we defined D14 as a functionally mature “young” state and D49 as an “aged” state characterized by functional decline and structural remodeling.

**Figure 1.**
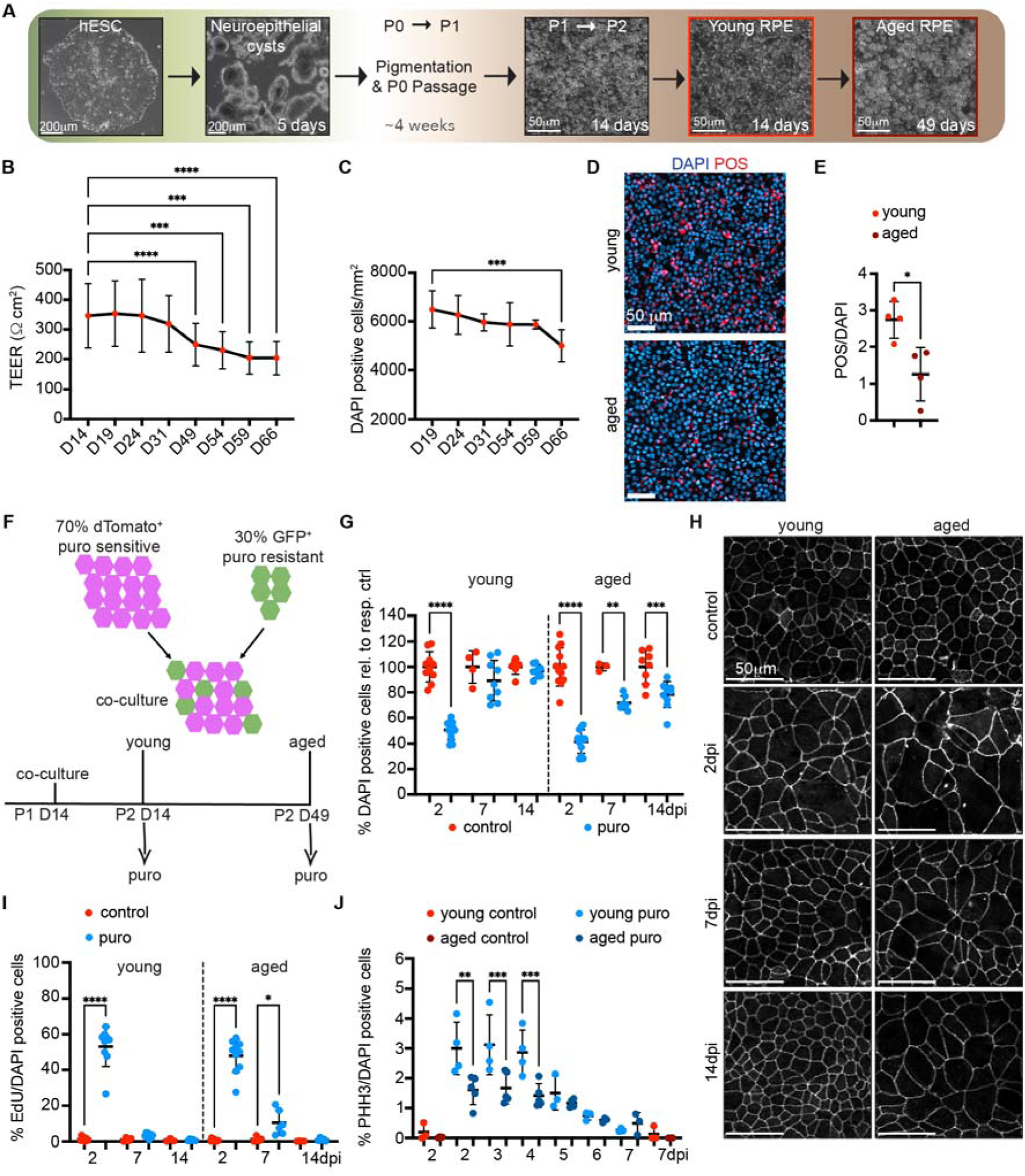
hESC-derived RPE recapitulate aging-associated dysfunction and impaired repair capacity *in vitro*. (A) hESC-RPE differentiation and extended culture schematic. (B and C) TEER (B; n = 21-32) and cell density (C; n = 3-12) after passage 2. (D and E) Representative images (D) and quantification (E; n = 4) of Alexa-Fluor 647-POS uptake. Scale bars, 50 μm. (F) Mosaic co-culture injury assay schematic. (G and H) Cell density after injury relative to age-matched controls (G; n = 3-13) and representative ZO-1 images (H). Scale bars, 50 μm. (I and J) Quantification of EdU^+^ S-phase cells (I; n = 3-10) and PHH3^+^ mitotic cells (J; n = 3-5) after injury. Data are mean ± SD. *n* represents independent RPE preparations. Mixed-effects model with Dunnett’s test (B), one-way ANOVA with Dunnett’s test (C), Welch’s t test (E), and two-way mixed-effects model with Šídák’s test (G, I, and J). \*\*\*\**p* < 0.0001, \*\*\**p* < 0.001, \*\**p* < 0.01, \**p* < 0.05. TEER, transepithelial electrical resistance; POS, photoreceptor outer segments; dpi, days post-injury. See also Figure S1.

Prolonged culture recapitulated several features of aged RPE. Transepithelial electrical resistance (TEER), a measure of epithelial barrier integrity, remained stable through D24 but declined thereafter, reaching a sustained reduction by D49 (Figure 1B). Despite barrier decline, D49 monolayers retained cobblestone morphology, ZO-1-positive tight junctions, and robust expression of the mature RPE marker Bestrophin-1 (BEST1) (Figure S1C), indicating preserved RPE identity. In parallel, cell density decreased (Figure 1C and S1C), while cellular and nuclear area increased (Figures S1A-S1C), consistent with age-associated RPE cell loss and compensatory enlargement observed *in vivo*^10,11^.

Prolonged culture also impaired specialized RPE functions. Phagocytosis of photoreceptor outer segments (POS), a critical homeostatic function that declines with age *in vivo*^12^, was reduced in D49 monolayers compared with D14 controls (Figures 1D and 1E). D49 cultures also showed delayed scratch-wound closure and larger residual wound areas (Figures S1D and S1E). Thus, this longitudinal hESC-RPE model captures key hallmarks of RPE aging, including barrier decline, reduced cell density with compensatory hypertrophy, impaired phagocytosis, and delayed wound closure.

### Aged RPE exhibit impaired injury-responsive repopulation and compensatory hypertrophy

Adult human RPE is predominantly quiescent in situ and remains largely postmitotic for decades^9,30^. Consistently, young and aged hESC-RPE monolayers showed minimal proliferative activity under homeostatic conditions (Figure S1F). To test whether localized cell loss triggers cell-cycle re-entry and repopulation, we developed a mosaic co-culture assay in which puromycin ablation produced a reproducible reduction in monolayer coverage (Figures 1F, S1G, and S1H). This enabled assessment of age-dependent injury-triggered cell-cycle re-entry and epithelial repair.

We first monitored epithelial barrier recovery. Both young and aged cultures exhibited a rapid TEER decline immediately after ablation. Young monolayers recovered to control levels within 7 days, whereas aged cultures required 14 days to return to their age-matched baseline (Figure S1I), reflecting delayed restoration to an already impaired state.

To determine whether barrier recovery reflected repopulation, we quantified cell density over the same time course. Young cultures largely recovered cell density by day 7, whereas aged cultures showed incomplete recovery even by day 14 (Figure 1G). Consistently, ZO-1 staining revealed persistent cellular hypertrophy in aged monolayers (Figures 1H and S1J), while DAPI staining indicated increased nuclear area (Figure S1K). This repopulation deficit prompted us to test whether aged RPE cells fail to enter the cell cycle after injury or instead enter S-phase but fail to progress to mitosis.

We therefore assessed EdU incorporation as a marker of S-phase entry and phospho-histone H3 (PHH3) as a marker of mitosis. Both young and aged cultures exhibited comparable EdU induction at 2 days post-injury (dpi), indicating that aged RPE cells retain the capacity for injury-induced S-phase entry (Figures 1I and S1L). However, aged RPE showed reduced mitotic progression, with fewer PHH3-positive cells during the peak proliferative window at 2-4 dpi (Figures 1J and S1M). Whereas EdU incorporation in young cultures returned to baseline by 7 dpi, aged RPE maintained elevated EdU levels (Figures 1I and S1L). Together with reduced PHH3 induction and incomplete cell-density recovery, these data indicate that aged RPE cells initiate S-phase after injury but fail to generate efficient mitotic output.

### Transcriptomic remodeling in hESC-RPE mirrors key features of human RPE aging *in vivo*

We performed bulk RNA sequencing on young and aged hESC-RPE to test whether our model recapitulates age-associated transcriptional changes observed in human RPE *in vivo*. Differential expression analysis identified 550 upregulated and 638 downregulated genes in aged versus young cultures (Table S1). Using gene set enrichment analysis (GSEA) with a published human donor RPE aging signature covering ages 31–93 years (Figure 2A)^31^, we found that aged hESC-RPE recapitulate age-associated transcriptional signatures observed in human RPE *in vivo*: genes downregulated with age *in vivo* showed concordant reduction in aged hESC-RPE (Figure 2B), whereas genes upregulated with age *in vivo* were concordantly enriched (Figure 2C). Together, these results support the use of aged hESC-RPE as a model of human RPE aging at the transcriptomic level.

**Figure 2.**
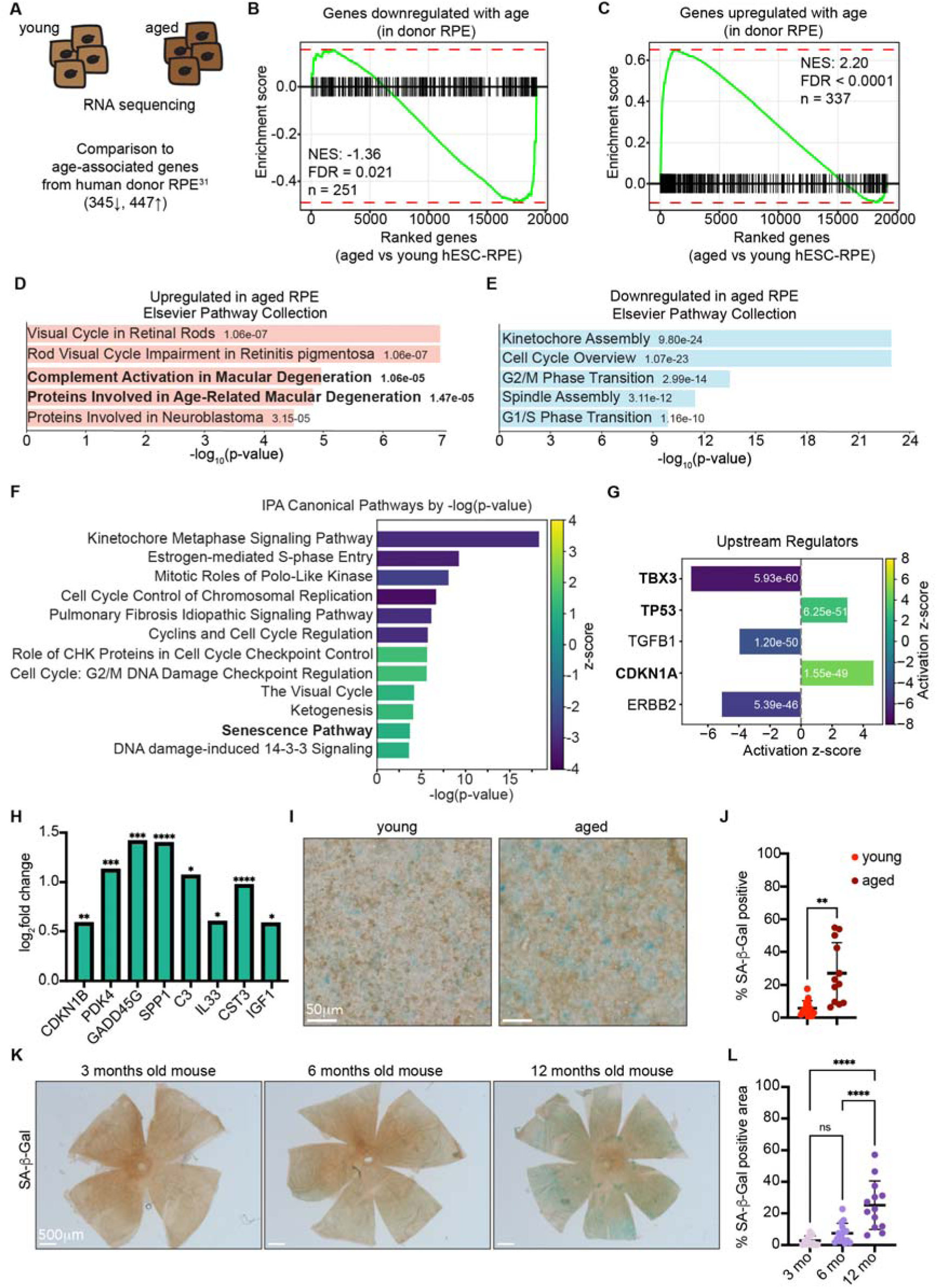
Aged hESC-RPE cells exhibit transcriptomic and cellular features of senescence. (A) GSEA comparison of aged hESC-RPE with human donor RPE signatures^31^. (B and C) GSEA showing enrichment of donor RPE age-downregulated (B) or age-upregulated (C) genes in aged hESC-RPE. (D and E) Elsevier Pathway Collection analysis of genes upregulated (D) and downregulated (E) in aged hESC-RPE. (F and G) IPA canonical pathways (F) and predicted upstream regulators (G) in aged versus young RPE. (I) Log_2_ fold change of selected senescence-associated genes. (I and J) Representative images (I) and quantification (J; n = 12-13) of SA-β-Gal^+^ cells. Scale bars, 50 μm. (K and L) Representative images (K) and quantification (L; n = 12-15 mice) of SA-β-Gal^+^ area in mouse RPE flatmounts. Scale bars, 500 µm. Data are mean ± SD. *n* represents independent RPE preparations (J) or individual mice (L); RNA-seq, n = 4 independent RPE preparations (B-H). Differential expression was performed using DESeq2 with Benjamini-Hochberg correction. Welch’s t test (J) and one-way ANOVA with Tukey’s test (L). \*\*\*\**p* < 0.0001, \*\*\**p* < 0.001, \*\**p* < 0.01, \**p* < 0.05. NES, normalized enrichment score; FDR, false discovery rate; IPA, Ingenuity Pathway Analysis; SA-β-Gal, senescence-associated β-galactosidase. See also Table S1.

Pathway enrichment of our data further revealed disease-relevant and immune remodeling in aged RPE, including increased AMD- and complement-associated pathways (Figure 2D). In contrast, young cultures were enriched for cell-cycle and mitotic programs, including kinetochore assembly and G2/M phase transition (Figure 2E). Ingenuity Pathway Analysis (IPA) yielded a similar pattern, with stronger representation of cell-cycle and mitotic programs in young RPE and enrichment of checkpoint-related programs in aged cultures, such as G2/M DNA damage checkpoint regulation and CHK-mediated cell-cycle control (Figure 2F). Because both states showed minimal baseline proliferation (Figure S1F), these differences are unlikely to reflect homeostatic proliferation. Rather, they suggest that young RPE retain a transcriptional state permissive for injury-induced cell-cycle progression, whereas aged RPE engage checkpoint-related programs that may restrict progression through mitosis. Together with reduced PHH3 induction after injury (Figures 1J and S1M), these results indicate that aged RPE acquire a senescence-associated state limiting cell-cycle completion after injury.

### Aged hESC-RPE exhibit features of cellular senescence

To identify regulators associated with age-related remodeling, we performed IPA upstream regulator analysis. This revealed predicted inhibition of the senescence-suppressive factor TBX3^32–34^ and activation of the pro-senescence regulators TP53 and CDKN1A (p21) (Figure 2G). Consistently, the cellular senescence pathway was enriched in aged cultures (Figure 2F), together with upregulation of the cell-cycle inhibitor *CDKN1B* (p27), the G2/M checkpoint regulator *GADD45G*, and the stress-associated metabolic regulator *PDK4* (Figure 2H).

Aged RPE also showed increased expression of secreted or extracellular factors, including *SPP1*, *C3*, *IL33*, *CST3*, and *IGF1* (Figure 2H). Together with complement pathway enrichment (Figure 2D), these changes are consistent with a SASP-like transcriptional program^35,36^. Notably, several of these factors, including *C3*, *IL33*, and *SPP1*, have also been implicated in AMD pathogenesis^37–39^. Thus, aged hESC-RPE acquires a senescence-associated transcriptional state marked by checkpoint-related programs and SASP-associated inflammatory and extracellular remodeling factors.

To validate the transcriptomic evidence of senescence at the cellular level, we stained for senescence-associated β-galactosidase (SA-β-Gal). Aged RPE monolayers contained a higher fraction of SA-β-Gal-positive cells, with ∼27% staining positive versus ∼6% in young controls (Figures 2I and 2J). Although these cells comprised a minority of the population, their accumulation paralleled the functional and transcriptional changes observed in aged cultures.

To assess whether cellular senescence also increases in RPE *in vivo*, we examined mouse RPE from 3 to 12 months of age. SA-β-Gal-positive RPE increased progressively, with pronounced accumulation by 12 months (Figures 2K and 2L). These findings indicate that aged hESC-RPE acquire senescence-associated transcriptional and cellular features, and SA-β-Gal accumulation *in vivo* supports the physiological relevance of this model.

### Age-associated repair impairment is conserved across independent hESC lines

We next tested whether these age-associated phenotypes were reproducible in an independent hESC line. H1-derived RPE showed similar cell-density decline and compensatory hypertrophy, whereas SA-β-Gal positivity trended upward in aged cultures but did not reach statistical significance (Figures S2A-S2D). Some functional readouts differed between lines: H1-derived RPE showed lower phagocytic activity that remained stable over time, whereas TEER declined in H9-derived RPE but remained relatively stable in H1-derived RPE (S2E-S2G). Despite these line-specific differences, impaired wound closure was conserved across both hESC-RPE lines (Figures S1D, S1E, S2H, and S2I). In the mosaic-injury assay, aged H1-derived RPE also showed a comparable repopulation deficit with persistent cellular hypertrophy (Figures S2J-S2M). Thus, despite line-specific differences, impaired injury-responsive repair is conserved across independent hESC-derived RPE models.

### Senolytic treatment reduces senescent cell burden in aged RPE

Senescent cells contribute to tissue dysfunction and impaired repair during aging and age-related diseases^40,41^, and senolytic strategies have emerged as potential therapeutic approaches^42,43^. Given the accumulation of senescence-associated features and conserved repair impairment across our aged RPE models, we hypothesized that reducing senescent cells might mitigate these phenotypes.

We screened established senolytics, including Navitoclax, Venetoclax, and Fisetin, in aged H9-derived RPE. Only the BCL-2/BCL-xL inhibitor Navitoclax significantly reduced cell density (Figures S3A and S3B). We therefore quantified the effect of Navitoclax on the SA-β-Gal-positive fraction in aged cultures (Figure 3A). Navitoclax reduced the SA-β-Gal-positive fraction dose-dependently (Figures 3B and S3C). At the lowest dose, SA-β-Gal-positive cells decreased by ∼41% relative to DMSO, while total cell density declined by only ∼15% (Figures 3B, 3C, and S3C), suggesting preferential senescent-cell depletion. This was associated with increased cleaved caspase-3 (CC3)-positive cells, consistent with apoptotic cell loss, and compensatory spreading by surviving cells (Figures S3D-S3G).

**Figure 3.**
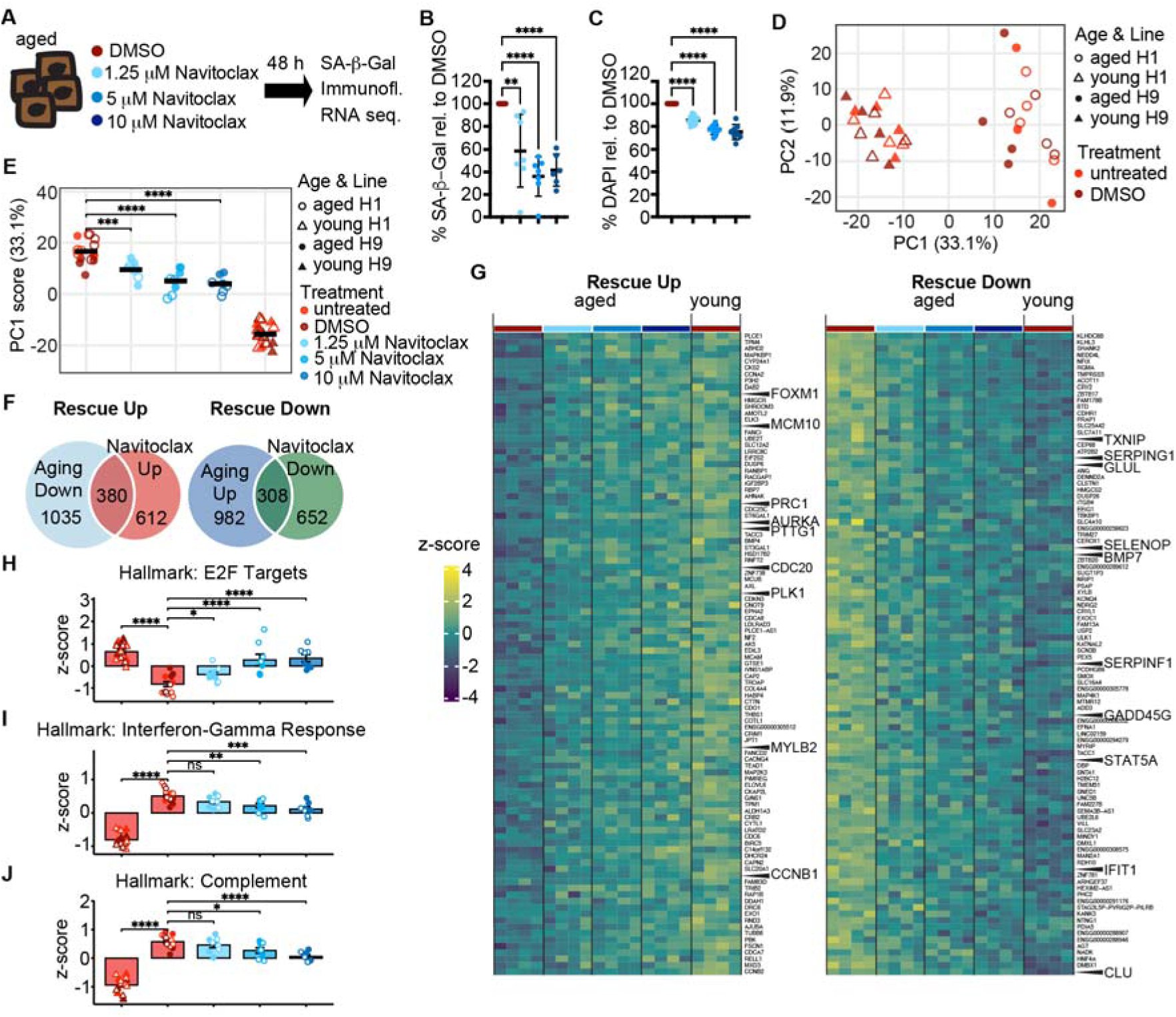
Senolytic treatment partially shifts aged hESC-RPE toward a younger transcriptional state. (A) Navitoclax treatment workflow. (B and C) Quantification of SA-β-Gal^+^ cells (B; n = 5-7) and DAPI^+^ nuclei (C; n = 8-12) after Navitoclax, normalized to DMSO. (D and E) PCA of H1- and H9-derived RPE transcriptomes defining the aging-associated PC1 axis (D), and PC1 scores across control and Navitoclax groups (E; n = 8 per treatment; n = 15-16 pooled controls). (F) Overlap between aging-associated and Navitoclax-responsive genes. (G) Heatmaps of top 100 Rescue-Up and Rescue-Down genes, with representative cell-cycle and inflammation/stress genes indicated. (H-J) Gene set scores for Hallmark E2F targets (H), Interferon-γ Response (I), and Complement (J). Data are mean ± SD (B-C) or mean ± SEM (H–J). *n* represents independent RPE preparations. One-way ANOVA with Dunnett’s tests (B and C), Wilcoxon rank-sum tests comparing each treatment to aged DMSO control (E), and linear mixed models with cell line as a random effect (H–J). Differential expression was performed using DESeq2 with Benjamini-Hochberg correction. \*\*\*\**p* < 0.0001, \*\*\**p* < 0.001, \*\**p* < 0.01, \**p* < 0.05; ns, not significant. PCA, principal component analysis. See also Figures S3 and S4 and Table S2.

In H1-derived RPE, Navitoclax similarly reduced SA-β-Gal-positive cells and total cell density despite lower baseline senescence burden (Figures S2C, S2D, and S3H-S3J), supporting reproducible senolytic effects across independent hESC-RPE models.

### Senolytic treatment shifts the aged RPE population transcriptome toward a younger state

To assess the effects of senolytic treatment, we performed RNA-seq on young, aged, and Navitoclax-treated aged RPE from H9- and H1-derived lines. Principal component analysis (PCA) revealed a conserved age-dependent transcriptomic axis separating young and aged cultures along PC1, which explained 33.1% of the variance (Figure 3D).

Consistent with our prior analysis (Figure 2D-F), genes contributing most strongly to the aging axis were enriched for inflammatory pathways, including Interferon-Gamma Response and Complement, alongside reduced mitotic programs (Table S2). Navitoclax shifted aged RPE dose-dependently toward the young transcriptomic state along PC1 (Figure 3E). At 10 μM, Navitoclax-treated aged cultures moved 39.9% toward the young baseline along PC1.

Differential expression analysis showed that, across all doses, Navitoclax induced gene expression changes that opposed age-associated changes (Figures S4A-S4C). We defined a Rescue signature of age-dysregulated genes shifted toward young levels after Navitoclax treatment, identifying 380 Rescue Up and 308 Rescue Down genes (Figure 3F). Across both lines, top-ranked Rescue genes shifted dose-dependently toward younger expression patterns (Figures 3G and S4D). Rescue Up genes included mitotic regulators such as *FOXM1*, *AURKA*, *CCNB1*, *PLK1,* and *CDC20*, whereas Rescue Down genes included inflammatory and stress-response genes, including *SERPING1*, *GADD45G*, *CLU*, *IFIT1,* and *SERPINF1*.

At the pathway level, Navitoclax partially restored E2F target gene expression while suppressing Interferon-Gamma Response and Complement programs (Figures 3H-3J and S4E). Together, these results indicate that Navitoclax shifts aged RPE toward a younger population-level transcriptomic state, partially restoring mitotic programs while suppressing inflammatory pathways, consistent with preferential depletion of senescence-associated cells.

### Senolytic priming enables structural repair and functional recovery following injury

Having established that Navitoclax shifts aged RPE toward a younger transcriptomic state, we asked whether it enhances injury-responsive repair in aged H9-derived RPE during a 21-day assay (Figure 4A). We selected the lowest effective dose, 1.25 µM, for all subsequent long-term assays. After a 48 h Navitoclax pulse, aged RPE maintained TEER comparable to DMSO controls for 17 days, indicating preserved barrier integrity despite reduced cell density (Figure 4B). However, Navitoclax alone did not induce repopulation (Figure 4C), indicating that the transcriptomic shift is insufficient to drive cell division without injury cues. The surviving RPE remained largely quiescent, with Ki67 positivity near or below 1% throughout the time course (Figure 4D), and compensated through progressive hypertrophy (Figure S4F).

**Figure 4.**
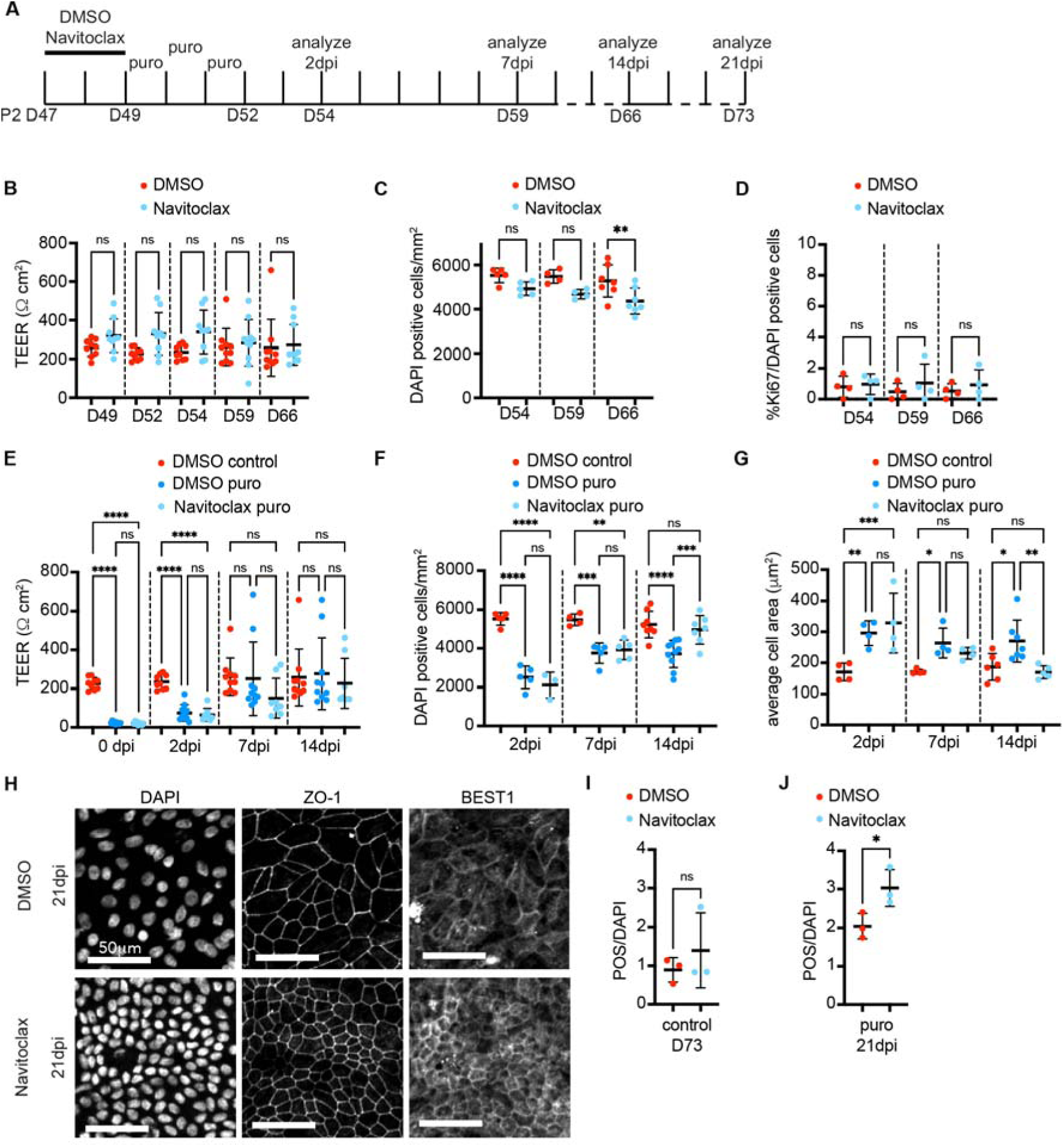
Senolytic priming enables injury-responsive repair and improves function in aged hESC-RPE. (A) Experimental timeline for Navitoclax (1.25 μM) pretreatment, mosaic injury, and longitudinal analysis. (B-D) TEER (B; n = 9-10), cell density (C; n = 4-7), and Ki67^+^ nuclei (D; n = 4) in uninjured aged H9-RPE treated with DMSO or Navitoclax. (E-G) TEER (E; n = 8-10), cell density (F; n = 4-10), and cell area (G; n = 4-7) after injury. (H) Representative DAPI, ZO-1, and BEST1 images at 21 dpi. Scale bars, 50 μm. (I and J) POS phagocytosis without injury (I; n = 3) or after injury (J; n = 3). Data are mean ± SD. *n* represents independent RPE preparations. Two-way mixed-effects model with Šídák’s test (B and E), two-way ANOVA with Šídák’s test (C and D), two-way ANOVA with Tukey’s test (F and G), and unpaired two-sided t test (I and J). \*\*\*\**p* < 0.0001, \*\*\**p* < 0.001, \*\**p* < 0.01, \**p* < 0.05; ns, not significant. See also Figure S4.

We therefore tested whether injury provided the additional cue required for productive repair. Using the mosaic injury assay (Figure 4A), TEER recovered toward baseline in both conditions (Figure 4E). However, DMSO-treated aged RPE compensated mainly through persistent hypertrophy, whereas Navitoclax-treated cultures showed substantial recovery of epithelial cell density and reduced cell area by 14 dpi (Figures 4F, 4G, and S4G). Thus, while neither senolytic treatment nor injury alone restored aged RPE cell density, their combination primed aged RPE for productive injury-responsive repopulation, a response we term senolytic priming.

To assess whether repopulated cultures retained RPE function, we examined BEST1 expression and phagocytosis at 21 dpi. Navitoclax-primed RPE formed dense, organized BEST1-positive monolayers, indicating retained RPE identity (Figure 4H). In uninjured cultures, Navitoclax alone did not improve phagocytosis (Figures 4I and S4H). However, after injury, Navitoclax-primed cultures showed enhanced POS uptake compared with injured DMSO controls, reaching levels comparable to young RPE (Figures 4J and S4I). Together, these findings show that senolytic treatment alone is insufficient to drive epithelial repair but primes aged RPE for injury-responsive repopulation and improved homeostatic function.

## DISCUSSION

In this study, we establish an hESC-RPE aging and injury model that reveals latent repair capacity in human RPE and shows that this capacity is suppressed with age. This decline in repair competence is associated with impaired mitotic progression, consistent with a senescence-associated G2/M impediment. Senolytic treatment alone is insufficient to drive epithelial repopulation, but primes aged RPE to respond productively to injury cues. Thus, reducing senescent cell burden creates a permissive state in which aged human RPE can re-engage injury-induced repair programs.

Our hESC-RPE aging model recapitulates key transcriptomic and functional features of aged human donor RPE, including inflammatory remodeling, reduced cell-cycle programs, functional decline, and senescent-cell accumulation. Previous long-term culture systems have modeled RPE wounding responses in fetal cells subjected to repetitive electrical ablation^44^ or transcriptomic maturation in hESC-RPE cultures^45^, but did not address how progressive cellular aging alters repair competence. Recent iPSC-RPE work showed that modest, apoptosis-driven density reductions in short-term cultures induced aging-like structural remodeling and impaired POS internalization through altered mechanical homeostasis^46^. Consistently, our extended cultures exhibit cell-density decline, compensatory hypertrophy, and functional impairment. By combining longitudinal aging with controlled mosaic ablation, our model distinguishes structural consequences of cell loss from age-dependent mechanisms limiting repopulation and identifies impaired mitotic progression as a barrier to RPE repair.

A key mechanistic insight is that aged RPE cells exhibit impaired mitotic progression rather than failed cell-cycle re-entry. Following injury, these cells incorporate EdU comparably to young cells but show fewer PHH3-positive cells, indicating S-phase entry with inefficient mitotic progression. Transcriptomic analyses support this interpretation, revealing suppression of mitotic programs alongside activation of checkpoint-related pathways, including CHK-mediated cell-cycle control and G2/M DNA damage checkpoint regulation. Together, these findings argue against complete cell-cycle incompetence and instead suggest that aged RPE retain proliferative machinery in a constrained state. This is consistent with observations in other quiescent tissues, including geriatric muscle satellite cells that transition from reversible quiescence to senescence-associated proliferative arrest^47^. More broadly, our findings support a model in which age-related repair failure reflects suppression of latent repair programs rather than irreversible loss of reparative potential.

Recent studies have shown that senolytics can reduce senescent RPE burden and improve retinal outcomes in acute stress-induced or genetic disease models^25–27^. We extend these findings by showing that, during gradual aging, senescent-cell accumulation contributes to impaired injury-responsive repair that can be relieved through senolytic priming. In our system, Navitoclax partially reverses age-associated gene expression at the population level, supporting the idea that a senescent subpopulation contributes to the inflammatory, mitotically constrained state of aged RPE. This pattern likely reflects preferential depletion of senescent cells rather than broad transcriptional reprogramming, a distinction requiring single-cell resolution. Importantly, functional recovery occurred only when senolytic treatment was followed by injury, suggesting that senolytic depletion removes a barrier to repair but does not itself provide the cues needed to initiate epithelial repopulation. Although enrichment for cells with higher proliferative capacity may contribute, the absence of spontaneous proliferation after Navitoclax alone, together with restored phagocytic function only after combined treatment, supports senolytic priming as a mechanism that re-enables injury-responsive repair rather than simply selecting proliferative clones.

This concept may extend beyond the RPE. Although senolytics show promise across aging and disease models, translation remains challenged by context-dependent efficacy, senescent cell heterogeneity, and incomplete clearance^48,49^. Our findings suggest that pairing senolytic priming with repair-activating stimuli may improve recovery in quiescent tissues where proliferation requires extrinsic cues. Consistent with this, BCL-2 family inhibition enhances age-impaired liver regeneration^50^, and topical Navitoclax pretreatment improves wound healing in aged skin^51^. Together with broader links between senescence and regenerative failure^52,53^, these studies support a framework in which endogenous repair programs persist in aged tissues but remain curtailed by senescence-associated constraints.

Several limitations warrant consideration. Although Navitoclax preferentially reduces senescent-cell burden, we cannot fully exclude off-target effects or broader population shifts. Our *in vitro* system also lacks key features of the retinal microenvironment, including Bruch’s membrane, systemic factors, immune components, and chronic environmental stressors. Moreover, our mosaic ablation assay enables controlled assessment of repair capacity but differs from the gradual dysfunction and regional atrophy of AMD. *In vivo*, adult human RPE primarily respond to damage through hypertrophy, migration, or transdifferentiation^4,5^. Whether the G2/M impediment identified here biases repair toward these compensatory responses rather than productive mitotic repopulation remains unknown. Future studies in AMD patient-derived iPSC-RPE and disease-relevant *in vivo* models, coupled with single-cell profiling, will be important for testing senolytic priming in pathological contexts.

Together, our findings identify cellular senescence as a targetable barrier to injury-responsive repair and introduce senolytic priming as a strategy for restoring latent repair competence in aged RPE. More broadly, they suggest that age-related repair failure may reflect active suppression of endogenous repair programs rather than irreversible loss of reparative potential. As senolytic therapies advance clinically, identifying safe repair-activating stimuli will be essential for improving tissue function in the aging eye.

## MATERIALS AND METHODS

### Cell lines and cell culture

Feeder-free hESC lines WA01 (H1, male) and WA09 (H9, female) were obtained from WiCell. Cells were regularly tested for mycoplasma contamination and confirmed negative. All hESCs were cultured on hESC-qualified Matrigel (Corning, 354277)-coated plates with mTeSR medium (STEMCELL Technologies, 85850) and maintained in a 5% CO_2_ incubator at 37°C. H9-GFPpuro (AAVS1-SA-2A-puro-2xCHS4-CAG-EGFP-WPRE-SV40-2xCHS4)^54^ and H9-dTom (AAVS1-CAGGS-dTomato-P2A-dTomato-WPRE)^55^ were gifts from J. Knoblich. To generate ubiquitously labeled, puromycin-resistant H1 cells, a reporter construct (AAVS1-SA-2A-puro-2xCHS4-CAG-dTomato-WPRE-SV40pA-2xCHS4)^56^ was inserted into the AAVS1 safe-harbor locus using TALEN technology as described^54,57^. Heterozygous insertion was verified by PCR.

RPE cells were differentiated as described previously^28,29^. Briefly, Dispase-dissociated hESC colonies were embedded in Matrigel (Corning, 354234) and cultured for 5 days in N2B27 medium. Following trypsinization of neuroepithelial cysts, 100,000-120,000 cells were plated on growth-factor-reduced Matrigel-coated 24-well Transwell inserts (Corning, 354230; Greiner, 662641) with 5 μM Y-27632 ROCK inhibitor for the first 24 h. Cells were maintained in RPE medium (DMEM + GlutaMAX, 20% KnockOut Serum Replacement, non-essential amino acids, 1 mM L-glutamine, penicillin-streptomycin, 0.1 mM β-mercaptoethanol) supplemented with 0.1 μg/ml Activin A (Qkine, Qk001). Medium was changed every 3 days for ∼4 weeks, until pigmentation and RPE morphology were observed. This stage was designated passage 0 (P0).

For passage 1 (P1), pigmented RPE were dissociated with 0.05% trypsin-EDTA (Gibco, 15400054) and seeded at 600,000 cells per 24-well Transwell insert or 1.8-2x10^6^ cells per 12-well Transwell insert (Greiner, 662641 or 665641). Cells were cultured for 2 weeks in RPE medium supplemented with 0.01 μg/ml Activin A for H9-derived RPE or without Activin A for H1-derived RPE, with medium changes every 3 days.

For passage 2 (P2), cells were dissociated with 0.05% trypsin-EDTA and seeded at 400,000 cells per 24-well Transwell insert (Greiner, 662641) or into 96-well glass-bottom plates (Greiner, 655891) in RPE medium without Activin A. Young and aged RPE were defined as P2 + 14 days (∼8 weeks total culture time) and P2 + 49 days (∼13 weeks total culture time), respectively.

### Transepithelial electrical resistance (TEER) measurements

TEER was measured using an EVOM^2^ epithelial volt/ohm meter with STX2 electrodes (World Precision Instruments). Measurements were performed after medium equilibration to room temperature. The resistance of blank inserts containing the same volume of medium was subtracted from each measurement. Values were then multiplied by membrane area (0.33 cm^2^) to calculate area-corrected TEER and are reported as Ω cm^2^.

### Immunofluorescence staining and imaging

Cells were fixed with 2% paraformaldehyde (PFA) for 30 min, washed with PBS, and treated with quenching solution (1X PBS, 100 mM glycine, 0.3% Triton X-100) for 20 min. Cells were blocked in blocking solution (1X PBS, 1% BSA, 0.3% Triton X-100) for 1 h at room temperature. Primary antibodies diluted in blocking solution were incubated overnight at 4°C. The following primary antibodies were used: ZO-1 (Invitrogen, 402200, 1:200), BEST1 (Abcam, ab2182, 1:500), phospho-histone H3 (PHH3; Novus, NB600-1168, 1:2000), Ki67 (BD Biosciences, 550609, 1:500), and cleaved caspase-3 (CC3; Cell Signaling, 9661S, 1:300). Alexa Fluor 488/568/647-conjugated secondary antibodies (Invitrogen) raised against the appropriate host species were used at 1:500 dilution. Nuclei were counterstained with DAPI (Sigma-Aldrich, D9542, 1:500) during the secondary antibody step. For EdU incorporation experiments, cells were incubated with 10 μM EdU for 24 h prior to fixation; EdU detection was performed using the Click-iT™ EdU Alexa Fluor 647 Imaging Kit (Invitrogen, C10340) according to the manufacturer’s protocol. Transwell insert membranes were excised, mounted in Mowiol (Carl Roth, 0713.1), and imaged on a Zeiss LSM 800 confocal microscope. All comparisons within an experiment were imaged using identical acquisition settings.

### Image analysis and quantification

Image analysis was performed using custom Python scripts utilizing Cellpose v2.2.3^58^ and scikit-image v0.24.0^59^. Cells were segmented from ZO-1 images using the Cellpose *cyto2* model and nuclei were segmented from DAPI images using the *nuclei* model. Cell area (μm^2^), nuclear area (μm^2^), and cell density were quantified using the *regionprops* function in scikit-image. For all conditions, five fields per well were imaged and analyzed. EdU- and Ki67-positive cells were identified using manual thresholds established from negative controls and applied consistently across experimental groups. CC3-positive cells were defined as cells with fluorescence intensity exceeding 3 standard deviations above background intensity.

### SA-β-Gal staining and analysis

SA-β-Gal staining was performed using the Senescence Cells Histochemical Staining Kit (Sigma-Aldrich, CS0030) according to the manufacturer’s instructions. Cultures were bleached with 5% H_2_O_2_ at 55°C to reduce pigmentation, counterstained with DAPI, mounted in Mowiol (Carl Roth, 0713.1), and imaged on a Zeiss Axio Imager Z2 microscope. Five fields per sample were imaged and analyzed across multiple independent experiments. Quantification was performed using an adapted Fiji macro based on Krzystyniak et al.^60^. Color thresholds were manually set to identify SA-β-Gal-positive area, and nuclei were identified by DAPI staining. Cells were scored as SA-β-Gal-positive if ≥30% of the nuclear area overlapped with SA-β-Gal staining; results are expressed as a percentage of total cells.

### Animal research and SA-β-Gal staining

Animals were maintained in accordance with Austrian animal welfare legislation and under institutional facility oversight. C57BL/6J mice were bred in-house and kept under specific pathogen-free conditions (22 ± 1°C; 55 ± 5% humidity; 12 h light/dark cycle). Male and female mice at 3, 6, and 12 months of age (n = 12-15 per group) were euthanized by CO_2_ asphyxiation. Eyes were immediately enucleated; the cornea, lens, and retina were removed, and eyecups were fixed for 7 min in fixation solution from the Senescence Cells Histochemical Staining Kit (Sigma-Aldrich, CS0030). SA-β-Gal staining was performed as described above. RPE flatmounts were bleached with 10% H_2_O_2_ at 55°C until pigmentation was sufficiently reduced for visualization, mounted in Mowiol, and imaged on a Zeiss Axio Zoom V16 stereomicroscope. Quantification was performed using a custom Fiji macro adapted from Krzystyniak et al.^60^. Images were converted to HSB color space, and thresholds were optimized per batch and applied uniformly to all samples within that batch to isolate SA-β-Gal signal. Results are expressed as the percentage of SA-β-Gal-positive area relative to total flatmount area. Both eyes from each animal were averaged to give a single value per mouse. Eyes with technical artifacts, such as incomplete bleaching, were excluded from analysis.

### Phagocytosis assay

Porcine photoreceptor outer segments (POS) were isolated as described previously^61^ and labeled with Alexa Fluor 647 (Thermo Scientific, A20006) for 2 h at 25°C with shaking at 500 rpm. Labeled POS were centrifuged at 9,000 x g for 10 min at 4°C, washed twice with wash buffer (10% sucrose, 20 mM phosphate buffer pH 7.2, 5 mM taurine), and resuspended in RPE medium. RPE monolayers were primed for 2 h with 5 μg/ml each of MFGE8 (R&D Systems, 2767-MF-050), GAS6 (R&D Systems, 885-GSB-050), and PROS1 (R&D Systems, 9489-PS-100), then incubated with labeled POS at ∼10 POS per cell for 24 h. Cells were washed five times with PBS containing 0.1 mM MgCl_2_ and 0.2 mM CaCl_2_ to remove unbound POS, fixed with 2% PFA for 30 min, and counterstained with DAPI. Transwell membranes were excised, mounted in Mowiol, and imaged on a Zeiss LSM 800. Five fields per sample were analyzed across multiple independent experiments. Confocal z-stacks spanning the full cell height were acquired, and maximum intensity projections were used for analysis. POS particles were scored as internalized when localized within the RPE cell volume in confocal z-stacks, and internalized Alexa Fluor 647-positive POS particles were quantified per cell using CellProfiler^62^. Nuclei were segmented by DAPI.

### Scratch assay

hESC-derived RPE monolayers in 96-well glass-bottom plates (Greiner, 655891) were scratched with a 10 μl pipette tip and imaged every 3 days for 14 days using a Zeiss Celldiscoverer 7. Scratch area was quantified using an adapted ImageJ plugin^63^ and expressed as percentage of total well area. Wells with initial scratch areas outside the predefined acceptable range were excluded from analysis to ensure uniform starting conditions.

### Senolytic drug treatment

Navitoclax (MedChemExpress, HY-10087), Venetoclax (MedChemExpress, HY-15531), and Fisetin (MedChemExpress, HY-N0182) were stored as 10 mM stocks in DMSO. For initial senolytic screening, aged H9-derived RPE were treated with Navitoclax, Venetoclax, or Fisetin at 30 nM, 100 nM, 300 nM, 1 μM or 3 μM, or with DMSO vehicle control, for 48 h. Based on screening results, Navitoclax was selected for detailed characterization. Aged RPE were treated with Navitoclax for 48 h at 1.25, 5, or 10 μM, with a 0.1% DMSO vehicle control matched to the highest drug concentration used. The lowest effective dose, 1.25 μM Navitoclax, was selected for all subsequent functional rescue experiments. For these experiments, cells were treated for 48 h with 1.25 μM Navitoclax or 0.0125% DMSO vehicle, after which the medium was replaced with standard RPE medium for the duration of the assay.

### RNA isolation and library preparation

Total RNA was extracted using TRI Reagent and purified with the Direct-zol RNA Miniprep Kit (Zymo Research, R2053). RNA integrity was assessed on a Fragment Analyzer (Agilent DNF 472 HS RNA Kit). For H9-derived aged versus young comparison (Figure 2), libraries were prepared from 500 ng total RNA using QuantSeq 3’ mRNA-Seq (Lexogen) and sequenced on an Illumina NovaSeq 6000 (S4 flow cell, XP workflow).

For the Navitoclax transcriptomic analysis (Figure 3), untreated and DMSO-treated young RPE, untreated and DMSO-treated aged RPE, and Navitoclax-treated aged RPE from H9-and H1-derived lines were prepared using Bulk RNA Barcoding and Sequencing (BRB-seq)^64^. BRB-seq libraries were prepared using 150 ng total RNA per sample to generate barcoded cDNA and amplified using a one-step RT/PCR reaction as described previously^64–66^. BRB-seq libraries were sequenced on an Illumina NovaSeq X (10B XP flow cell). All sequencing was performed in paired-end 150 bp mode.

### RNA-seq data analyses

Three to four independent biological replicates were used per condition.

QuantSeq reads were processed with the nf-core/rnaseq pipeline (v3.10)^67^ using default parameters. Quality control was performed with FastQC (v0.11.9). Reads were aligned to GRCh37 with STAR (v2.6.1d)^68^, and gene-level quantification was performed with Salmon (v1.9.0)^69^ in alignment mode. Unique molecular identifiers (UMIs; 6 bp) were extracted and deduplicated using UMI-tools (v1.1.2)^70^ to account for PCR bias. Gene counts were imported into R (v4.3.0), and differential gene expression analysis performed with DESeq2 (v1.40.2)^71^ using Benjamini-Hochberg FDR correction. Genes with FDR < 0.05 and |log₂ fold change| > 0.5 were considered significant. For comparison with human donor RPE aging signatures^31^, the published list of age-associated genes was tested for enrichment in aged versus young hESC-RPE using gene set enrichment analysis. Pathway and upstream regulator analyses were performed using Ingenuity Pathway Analysis (IPA; QIAGEN) and Enrichr^72,73^ against the Elsevier Pathway Collection.

BRB-seq reads were trimmed for adapters and polyA tails with Cutadapt (v1.18)^74^ and paired with SeqKit (v2.7.0)^75^. Reads were aligned to GRCh38, demultiplexed using sample barcodes, and quantified with STARsolo (v2.7.11a)^76^. Count matrices were loaded into R (v4.3.0), and differential expression analysis was performed with DESeq2 (v1.40.2)^71^ with age and treatment as covariates. Genes with fewer than 10 total counts were excluded. Variance-stabilizing transformation (VST) was applied for normalization. Cell line batch effects were removed using *removeBatchEffect* from limma v3.58.1^77^ while preserving age and treatment effects. Genes with FDR < 0.05 and |log_2_ fold change| > 0.5 were considered significant.

#### Principal component analysis and rescue quantification

PCA was performed on batch-corrected expression values from control samples, defined as young and aged untreated or DMSO-treated samples, using the top 5,000 most variable genes. Navitoclax-treated samples were then projected into this control PCA space to assess treatment-associated transcriptional shifts. PC1 scores were compared between prespecified groups using Wilcoxon rank-sum tests. PC1-based rescue toward a young transcriptional state was calculated as the shift in mean PC1 score from aged controls toward young controls, expressed as a percentage of the total PC1 distance between aged and young controls.

#### Gene set enrichment analysis

GSEA was performed on PC1 gene loadings using fgsea (v1.26.0)^78^ with MSigDB Hallmark gene sets retrieved via msigdbr (v25.1.1)^79^. Gene sets containing 15–500 genes were included and filtered at FDR < 0.05. Module scores for selected Hallmark pathways were calculated as the mean z-score of leading-edge genes. Statistical comparisons were performed using linear mixed models with cell line as a random effect using lme4 (v1.1.37) and lmerTest (v3.1.3).

#### Rescue gene identification

Rescue genes were defined as those significantly altered with aging (FDR < 0.05, |log₂ fold change| > 0.5) and significantly reversed by Navitoclax in the opposite direction. The degree of transcriptional reversal was assessed by Pearson correlation between aging-associated and Navitoclax-associated log₂ fold changes. Heatmaps were generated using ComplexHeatmap (v2.16.0).

### Statistical analysis

Statistical analyses were performed using GraphPad Prism (v10.6.1) and R (v4.3.0). No statistical methods were used to predetermine sample sizes. For all experiments, *n* represents independent biological replicates defined as independent RPE preparations for *in vitro* experiments and individual animals for *in vivo* experiments. Technical replicates or imaging fields of view were averaged within each biological replicate before statistical testing unless otherwise indicated. Data are presented as mean ± standard deviation (SD) unless otherwise indicated. Box plots display median, 25th–75th percentiles, and min–max values. For two-group comparisons, unpaired two-sided Student’s t tests or Welch’s t tests were used, with Welch’s correction applied for unequal variances. For multiple comparisons, one-way or two-way ANOVA was used with Dunnett’s test for comparisons to control, Tukey’s test for all pairwise comparisons, or Šídák’s test for selected pairwise comparisons, as appropriate. For longitudinal repeated-measures data, mixed-effects models were used with post hoc comparisons as indicated in figure legends. For non-normal data, Wilcoxon rank-sum tests were used. For RNA-seq GSEA across treatment groups, linear mixed models with cell line as a random effect were used to account for cell line effects. Statistical significance was defined as *p* < 0.05. Exact *p* values or significance levels are indicated in figure legends.

### Data availability

RNA-seq datasets generated in this study are being deposited in the NCBI Sequence Read Archive (SRA) and will be publicly available as of the date of publication. Source data are provided with the paper. Any additional information required to reanalyze the data reported in this paper is available from the lead contact upon request.

## ACKNOWLEDGMENTS

We thank the Tanaka laboratory for valuable discussions and support. We are especially grateful to Katharina Lust, Elad Bassat, and Thijs Brouwer for careful reading of the manuscript and helpful feedback. We thank Jiaye Yang, Stefanie Horer, Alexander Wöhrleitner, and Nicole Leeb for assistance with cell culture, and Salvador Gonzalez Juarez for assistance with QuantSeq data processing. We acknowledge the Next Generation Sequencing Facility of the Vienna BioCenter Core Facilities (VBCF) and the IMP/IMBA/GMI BioOptics Facility for expert technical assistance, specifically Pawel Pasierbek, Alberto Moreno Cencerrado, Gabriele Bradamante, and Thomas Lendl. We also thank the Vienna BioCenter animal facility staff for their continuous support. We thank Jürgen Knoblich for providing cell lines and plasmids, and Christopher Esk for discussions and technical support, and the IMBA Stem Cell Core Facility for the generation of hESC lines. We further thank Bill Keyes for insightful discussions and critical reading of the manuscript, and Martin Schönlein for careful reading of the manuscript. This work was funded by Boehringer Ingelheim Pharma GmbH & Co. KG. B.R. received funding from the European Union’s Framework Programme for Research and Innovation Horizon 2020 (2014-2020) under the Marie Curie Skłodowska Grant Agreement Nr. 847548. F.B.P. is supported by a DOC Fellowship of the Austrian Academy of Sciences at the University of Vienna. For the purpose of open access, the author has applied a CC BY public copyright licence to any Author Accepted Manuscript version arising from this submission.

## AUTHOR CONTRIBUTIONS

Conceptualization, B.R. and E.M.T.; Investigation, B.R., C.E., and F.B.P.; Image analysis, B.R., K.I., and C.E.; Bioinformatics analysis, B.R. and F.B.P.; Resources, S.A.; Writing – original draft, B.R.; Writing – review & editing, all authors; Funding Acquisition, E.M.T., H.N., and R.A.B.

## Declaration of generative AI and AI-assisted technologies in the manuscript preparation process

During the preparation of this work the authors used Abacus.AI ChatLLM tools to improve language and readability. After using these tools, the authors reviewed and edited the output as needed and take full responsibility for the content of the publication article.

---

**Figure S1.**
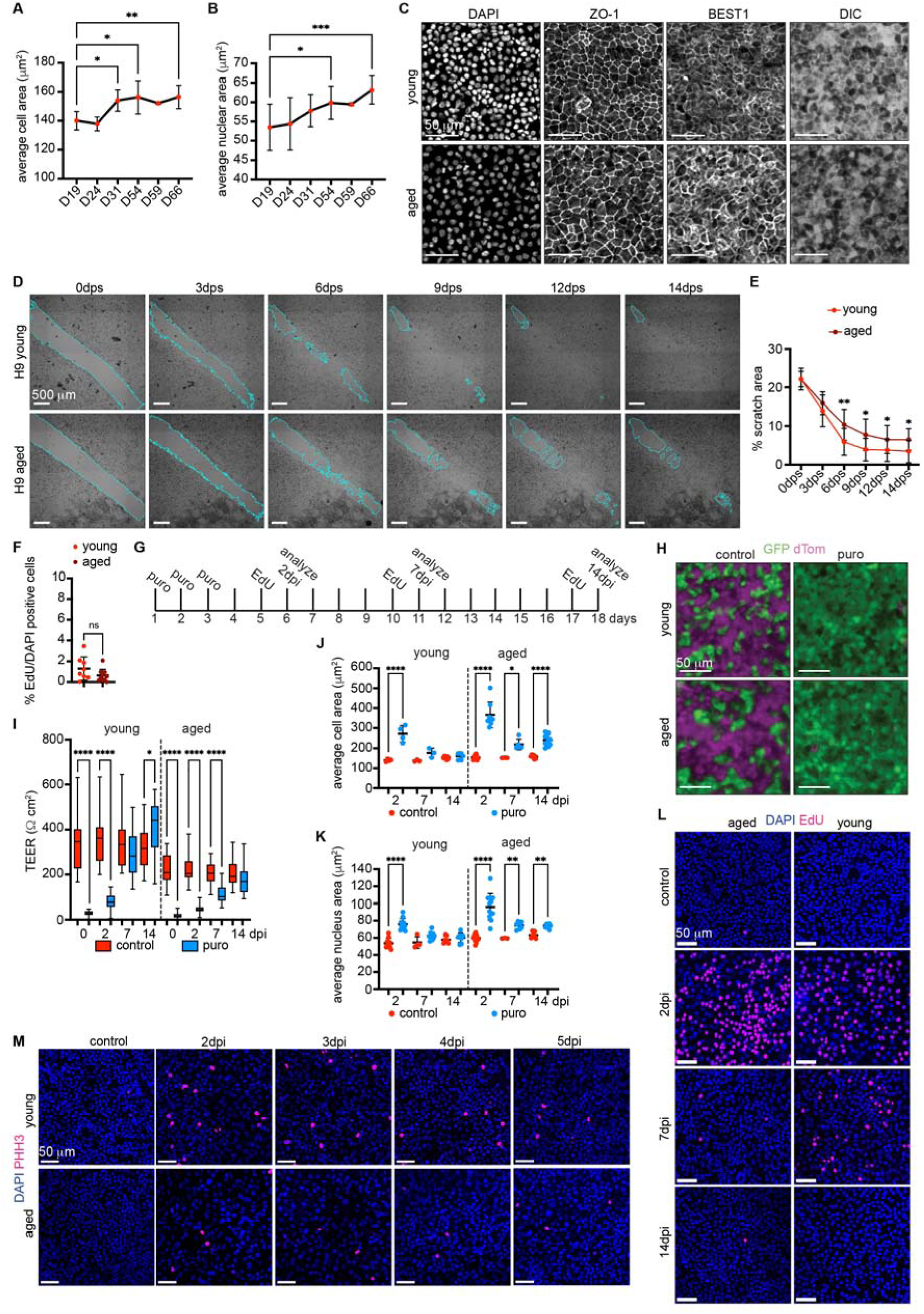
Establishment and characterization of the longitudinal hESC-RPE aging and mosaic injury model. Related to Figure 1. (A and B) Quantification of average cell area (A; n = 3-7) and nuclear area (B; n = 3-12) from D19-D66. (C) Representative images of young (D14) and aged (D49) hESC-RPE stained for DAPI, ZO-1, and BEST1, with corresponding DIC images. Scale bars, 50 μm. (D and E) Representative scratch images (D) and quantification of percentage scratch area over time (E; n = 11-15). Scratch area is outlined in cyan. Scale bars, 500 μm. (F) Baseline percentage of EdU^+^ cells in young and aged hESC-RPE under uninjured conditions (n = 8-10). (G) Experimental timeline of puromycin-induced mosaic injury and analysis at 2, 7, and 14 days post-injury (dpi). (H) Representative images of GFP+ (puromycin-resistant; green) and dTomato+ (puromycin-sensitive; magenta) co-cultures in control and puromycin-treated young and aged RPE. Scale bars, 50 μm. (I) TEER time course in control and injury groups for young and aged cultures (n = 21-30). (J and K) Quantification of average cell area (J; n = 3-9) and nuclear area (K; n = 3-14) after injury in young and aged RPE. (L) Representative DAPI (blue) and EdU (magenta) staining across the injury time course. Scale bars, 50 μm. (M) Representative DAPI (blue) and PHH3 (magenta) staining across 2-7 dpi. Scale bars, 50 μm. Data are mean ± SD. *n* represents independent RPE preparations. One-way ANOVA with Dunnett’s test (A and B), two-way mixed-effects model with Tukey’s test (E), Welch’s t test (F), mixed-effects model with Šídák’s test (I), and two-way ANOVA with Šídák’s test (J and K). \*\*\*\**p* < 0.0001, \*\*\**p* < 0.001, \*\**p* < 0.01, \**p* < 0.05. DIC, differential interference contrast; dps, days post-scratch; TEER, transepithelial electrical resistance; dpi, days post-injury.

**Figure S2.**
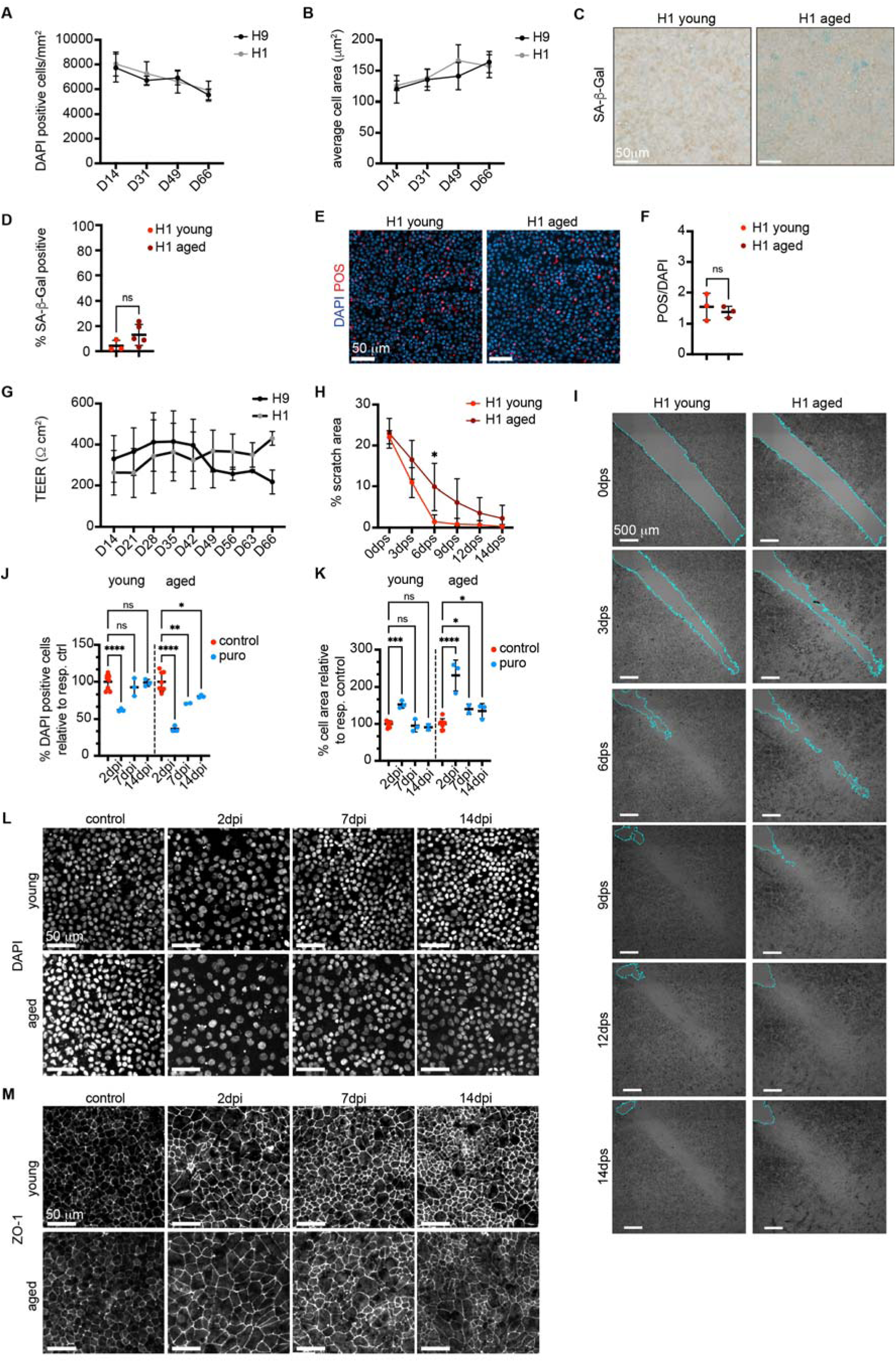
Aging-associated phenotypes are recapitulated across independent hESC-RPE lines. Related to Figures 1 and 2. (A and B) Cell density (A; n = 2-8) and average cell area (B; n = 3-6) over time in H9- (black) and H1-derived (gray) RPE. (C and D) Representative SA-β-Gal (C) staining and quantification of SA-β-Gal-positive cells as a percentage of DAPI^+^ nuclei (D; n = 3-5) in young and aged H1-derived RPE. Scale bars, 50 μm. (E and F) Representative images of POS uptake (E) and quantification of phagocytic capacity (F; n = 3) in young and aged H1-derived RPE. Scale bars, 50 μm. (G) TEER over time in H9- (black) and H1-derived (gray) RPE from D14-D66 (n = 3-18). (H and I) Quantification of scratch area over time (H; n = 3-5) and representative images (I) in H1-derived RPE. Scratch area is outlined in cyan. Scale bars, 500 μm. (J and K) Cell density (J; n = 2-8) and average cell area (K; n = 3-6) after puromycin injury in H1-derived RPE, normalized to age-matched controls. (L and M) Representative DAPI (L) and ZO-1 (M) images after injury in H1-derived RPE. Scale bars, 50 μm. Data are mean ± SD. *n* represents independent RPE preparations. Two-way ANOVA with Šídák’s test (A, B, J and K), Welch’s t test (D), unpaired two-sided t test (F), two-way mixed-effects model with Šídák’s test (G), and two-way mixed-effects model with Tukey’s test (H). \*\*\*\**p* < 0.0001, \*\*\**p* < 0.001, \*\**p* < 0.01, \**p* < 0.05; ns, not significant. POS, photoreceptor outer segments; TEER, transepithelial electrical resistance; dpi, days post-injury.

**Figure S3.**
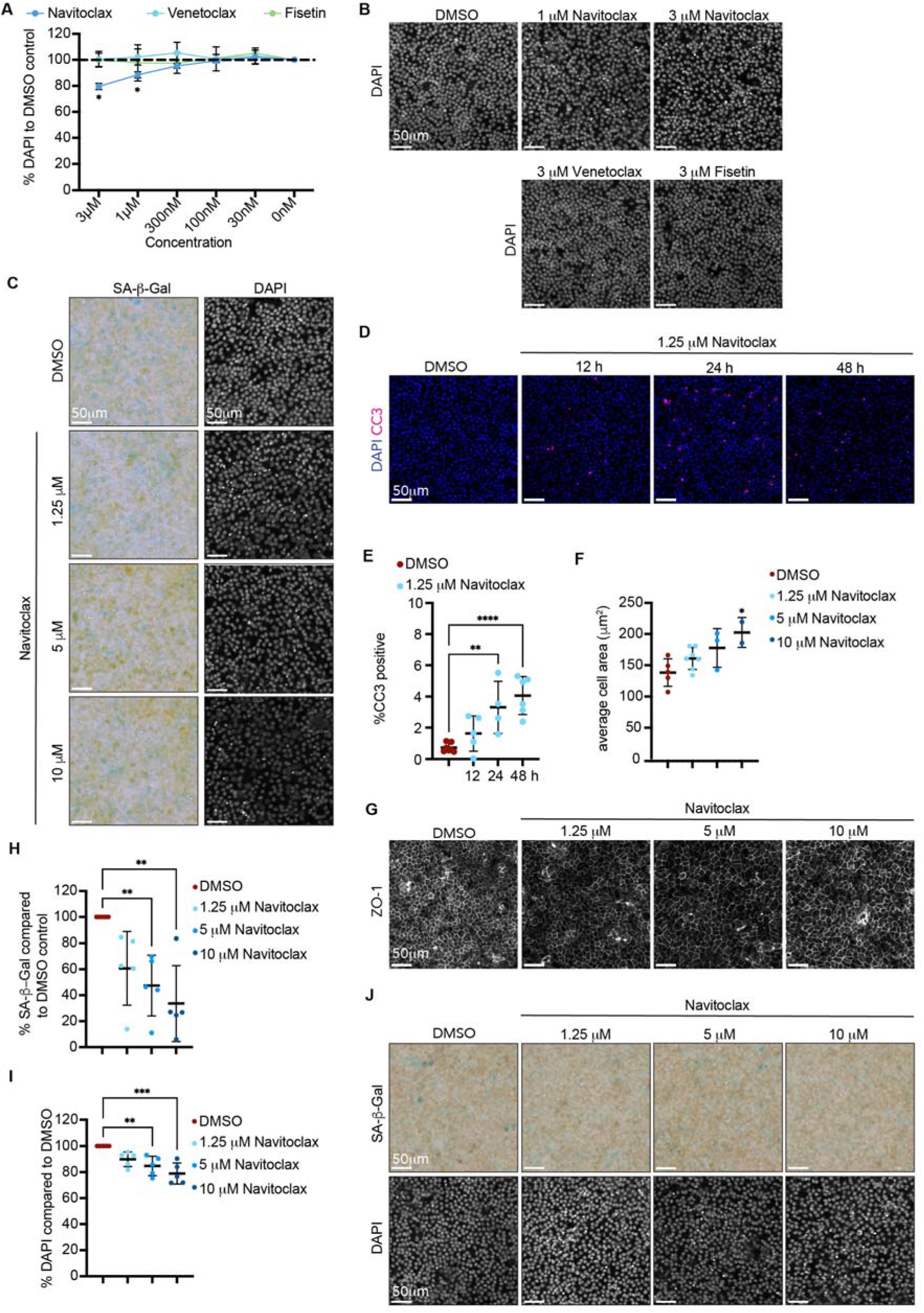
Validation of Navitoclax as a senolytic in hESC-RPE. Related to Figure 3. (A) Dose-response quantification of cell density in aged H9-derived RPE after 48 h treatment with Navitoclax, Venetoclax, or Fisetin, normalized to DMSO (n = 3). (B) Representative DAPI images of aged H9-derived RPE after treatment with the indicated compounds and concentrations. Scale bars, 50 µm. (C) Representative SA-β-Gal and DAPI staining in aged H9-derived RPE after treatment with the indicated Navitoclax doses. Scale bars, 50 µm. (D and E) Representative DAPI (blue) and cleaved caspase-3 (CC3; magenta) images (D) and quantification of CC3^+^ cells as a percentage of DAPI^+^ nuclei (E; n = 5-7) at indicated time points after 1.25 µM Navitoclax. Scale bars, 50 µm. (F and G) Quantification of average cell area (F; n = 2-7) and representative ZO-1 images (G) in aged H9-derived RPE after 48 h DMSO or Navitoclax treatment at the indicated doses. Scale bars, 50 µm. (H-J) Quantification of SA-β-Gal^+^ cells (H) and DAPI^+^ nuclei (I) in aged H1-derived RPE after 48 h Navitoclax treatment, normalized to DMSO (n = 5), and representative SA-β-Gal and DAPI images (J). Scale bars, 50 µm. Data are mean ± SD. *n* represents independent RPE preparations. Two-way mixed-effects model with Dunnett’s test (A), one-way ANOVA with Dunnett’s test (E and F), and one-way ANOVA with Šídák’s test (H-I). \*\*\*\**p* < 0.0001, \*\*\**p* < 0.001, \*\**p* < 0.01, \**p* < 0.05.

**Figure S4.**
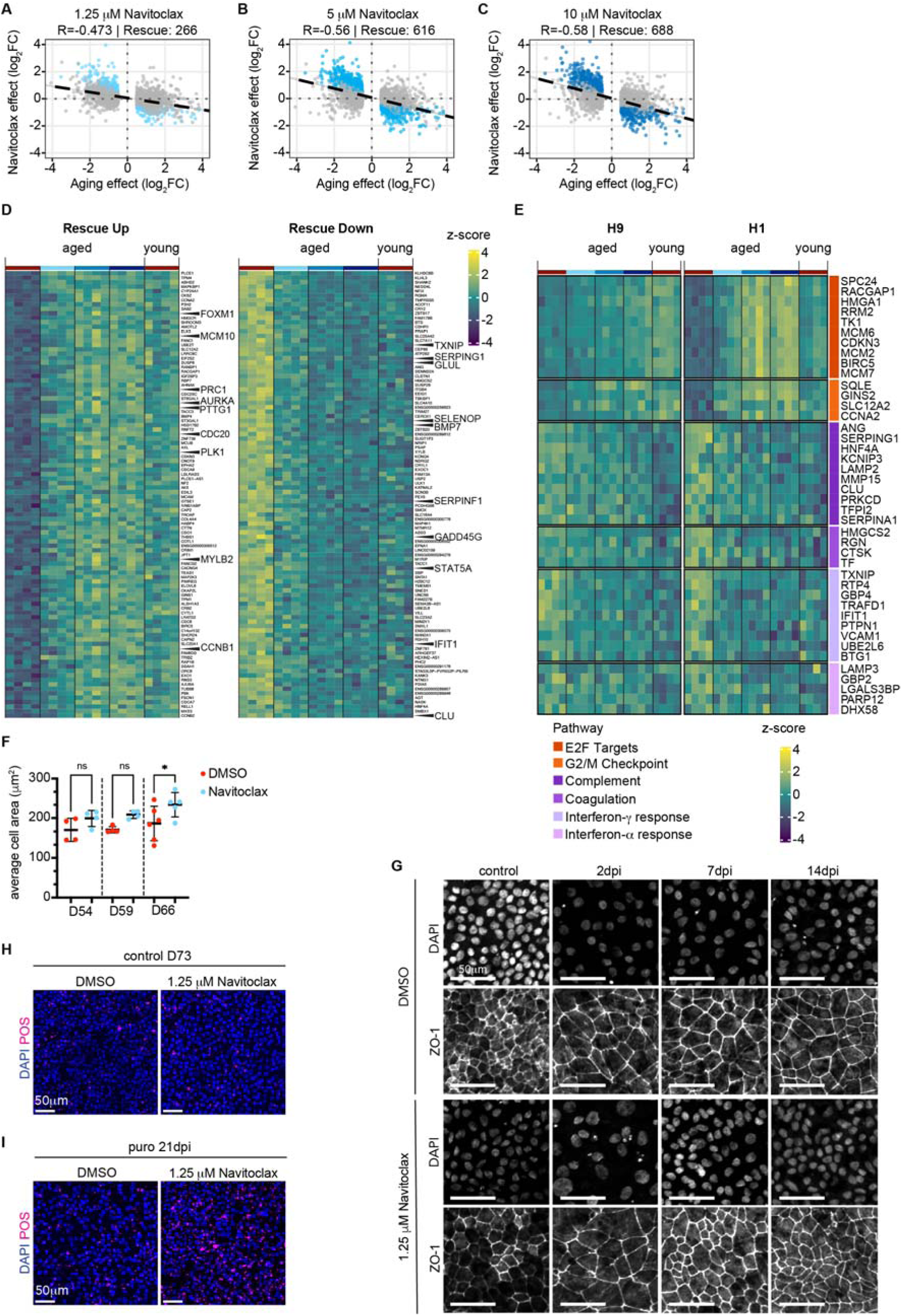
Transcriptional and functional characterization of Navitoclax responses in aged hESC-RPE. Related to Figures 3 and 4. (A-C) Scatter plots comparing aging-associated gene expression changes (aged DMSO versus young DMSO; *x* axis) with Navitoclax-induced changes in aged RPE (aged Navitoclax versus aged DMSO; *y* axis) for aging-associated differentially expressed genes (adjusted *p* < 0.05, |log_2_FC| > 0.5), in a combined, line-corrected model across H9- and H1-derived RPE with 1.25 µM (A; R = - 0.473, Rescue = 266), 5 µM (B; R = -0.56, Rescue = 616), and 10 µM (C; R = -0.58, Rescue = 688) Navitoclax. Pearson correlation coefficients (*R*) and the number of rescued genes are indicated; rescued genes were defined as genes showing opposite directional changes in aging and Navitoclax treatment (adjusted *p* < 0.05 and |log*_2_*FC| > 0.5) and are highlighted in blue. (D) Heatmaps of the top 100 Rescue-Up and top 100 Rescue-Down genes ranked by a hybrid score combining statistical significance and reversal magnitude; representative cell-cycle (up) and inflammation/stress (down) genes are labeled. (E) Heatmaps of leading-edge genes from selected Hallmark pathways (E2F targets, G2/M checkpoint, complement, coagulation, interferon-γ response, and interferon-α response) in H9- and H1-derived RPE, shown as z-scores. (F) Quantification of average cell area in aged H9-derived RPE after 48 h treatment with DMSO or 1.25 µM Navitoclax in the absence of injury at D54, D59, and D66 (n = 4-6). (G) Representative images of aged H9-derived RPE pretreated with DMSO or 1.25 µM Navitoclax, stained for DAPI and ZO-1 in uninjured and injured conditions at 2, 7, and 14 dpi. Scale bars, 50 µm. (H and I) Representative images of DAPI (blue) and POS (magenta) staining after DMSO or 1.25 µM Navitoclax pretreatment in uninjured control RPE (H) and injured RPE at 21 dpi (I). Scale bars, 50 µm. Data are mean ± SD. *n* represents independent RPE preparations. Two-way ANOVA with Šídák’s test (F). \**p* < 0.05, ns, not significant. RNA-seq differential expression analysis and gene ranking were performed as in Figure 3. POS, photoreceptor outer segments; dpi, days post-injury.

**Supplementary Table S1. Differentially expressed genes in aged versus young hESC-derived RPE (H9). Related to Figure 2**.

Bulk RNA-seq was performed on young and aged hESC-derived RPE monolayers (n = 4). Differential expression was analyzed with DESeq2. Genes with adjusted p < 0.05 and |log_2_ fold change| > 0.5 are included.

**Supplementary Table S2. Hallmark GSEA of genes contributing to PC1 (aging axis). Related to Figure 3**.

Hallmark GSEA of genes ranked by their contribution to PC1 (aging axis) in Figure 3D. Positive NES values indicate enrichment toward the aged direction of PC1, and negative NES values indicate enrichment toward the young direction. Pathways with FDR < 0.05 are shown.

